# A uniform benchmark for testing ssrA-derived degrons in the Escherichia coli ClpXP pathway

**DOI:** 10.1101/2021.08.26.457770

**Authors:** Maria Magdalena Klimecka, Anna Antosiewicz, Matylda Anna Izert, Patrycja Emanuela Szybowska, Piotr Krzysztof Twardowski, Clara Delaunay, Maria Wiktoria Górna

## Abstract

The ssrA degron is commonly used in fusion proteins to control protein stability in bacteria or as an interaction module. These applications often rely on the modular activities of the ssrA tag in binding to the SspB adaptor and in engaging the ClpXP protease. However, a comparison of these activities for a substantial standard set of degron variants has not been conducted previously, which may hinder developments of new variants optimized exclusively for one application. Here, we strive to establish a benchmark that will facilitate the comparison of ssrA variants under uniform conditions. In our workflow, we included methods for expression and purification of ClpX, ClpP, SspB and eGFP-degrons, assays of ClpX ATPase activity, of eGFP-degron binding to SspB and for measuring eGFP-degron degradation *in vitro* and *in vivo*. Using uniform, precise and sensitive methods under the same conditions on a range of eGFP-degrons allowed us to determine subtle differences in their properties that can affect their potential applications. Our findings can serve as a reference and a resource for developing targeted protein degradation approaches.

**SUMMARY:** This work lays standards for assays used to compare engagement of SspB and ClpXP by a set of ssrA-derived degrons that can be used to fine-tune tools for protein stability control.

## INTRODUCTION

A significant part of the directed protein degradation methods is based on the use of degrons. Knowledge of their properties, the strength of binding to the components of the degradation machinery or the impact on the efficiency of degradation is necessary for their skillful use in fusion proteins targeted for degradation or as interaction modules. One of the best described bacterial degrons which is the most frequently used in biotechnology is the ssrA tag sequence and its variants. In the bacterial quality control pathway, ssrA tag is added in a process called trans-translation. During trans-translation, tmRNA (transfer-messenger RNA) enters stalled ribosomes, replaces the mRNA and promotes the translation of a peptide tag (ssrA) encoded within tmRNA onto the C terminus of the nascent polypeptide (Keiler, 2008). The ssrA tag is conserved across bacteria and in *Escherichia coli* is recognized mainly by the ClpXP protease, which is a member of the ATPases Associated with diverse cellular Activities (AAA+) family. ClpX is the unfoldase subunit comprising a hexameric ring decorated with dimers of the ClpX N-terminal Zinc Binding Domain (ZBD), while the peptidase ClpP forms a double ring of heptamers that can bind ClpX hexamers on one or both faces of the ClpP barrel (Fig 1). The ssrA tag consists of 11 amino acids (AANDENYALAA) in which the AANDENY motif is responsible for binding of the ClpX adaptor, i.e. SspB protein, and the ALAA motif binds to the ClpX unfoldase central pore itself (Baker & Sauer, 2012). This modularity of ssrA is frequently exploited in biotechnology for various applications.

**Figure 1.**
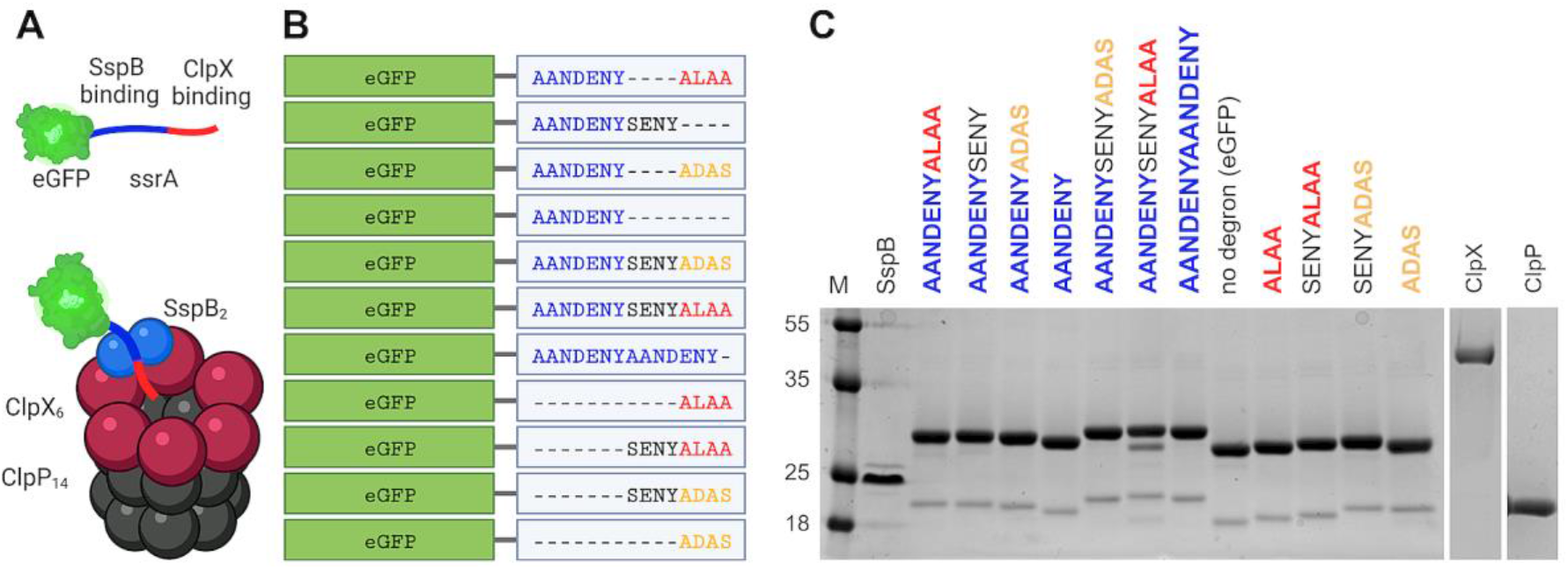
Characteristics of the studied proteins. (A) A schematic representation of the ClpXP protease machinery: the complex of the tetradecameric ClpP barrel-like structure (ClpP_14_; gray) and the homohexameric ClpX ring (ClpX_6_; purple), with an attached SspB dimer (SspB_2_; blue) facilitating the eGFP-degron (green) binding to ClpX. The figure was created with BioRender.com and Mol* (Sehnal *et al*, 2018). (B) Fusions of the ssrA tag and its modified variants with eGFP protein used in the study. (C) SDS-PAGE electrophoresis protein profiles of the SspB, eGFP-degrons, ClpX and ClpP proteins used in the study.

The biochemical parameters for various variants of the ssrA tag have already been described in the literature with the use of multiple experimental setups and diverse methods. However, typically only a few selected variants have been characterized per publication, making it difficult to properly compare those variants that have been described using different experimental setups. Moreover, there is no simultaneous comparison of the degradation tag properties, such as SspB binding and degradation efficiency, for a substantial set of ssrA variants.

Herein, we strive to establish a benchmark that will facilitate the comparison of the ssrA degron and its variants under uniform conditions. In our workflow, we included methods for the expression and purification of ClpX, ClpP, SspB, eGFP-degron proteins (fusions of eGFP with variants of the ssrA tag (Fig. 1A & 1B)), tests for the ATPase activity of ClpX, measurements of eGFP-degrons binding to the SspB protein and methods for measuring the degradation of eGFP-degrons *in vitro* and *in vivo*. Using uniform, precise and sensitive methods under the same conditions on a range of eGFP-degrons allowed us to determine subtle differences in their properties, their necessary and negligible elements, and to indicate their potential applications. Our findings can be a valuable point of reference and a resource for researchers using targeted protein degradation approaches.

## METHODS

### Cloning of clpP, clpX and sspB genes to pET28a expression vector by Sequence and Ligation Independent Cloning (SLIC)

DNA encoding full-length *clpP, clpX* and *sspB* genes of *Escherichia coli* (Top10 strain) was obtained by colony PCR amplification using specific primers (Suppl. Table 1) containing sequences complementary to pET28a plasmid with His tag and SUMO protease cleavage site.

The SLIC reaction was performed by mixing 100 ng of pET28a linearized empty vector, 100 ng of PCR-amplified DNA fragment (with *clpP, clpX* or *sspB* gene) and 1U of T4 DNA polymerase (Thermo Scientific) in 1X FastDigest Green Buffer (Thermo Scientific) in a volume of 30 µl. After 30 min incubation in RT, reactions were stopped by adding dCTP (EurX) to a final concentration of 1 mM, and the tubes were incubated for 30 minutes at 37°C for annealing. Finally, reactions were transformed to *E. coli* Top10 for selection on kanamycin.

For the preparation of pBAD-eGFP-degron constructs (Suppl. Table 2), we modified eGFP-pBAD plasmid (later referred to as pBAD-eGFP) containing His-TEV-tagged eGFP gene, obtained from Michael Davidson (Addgene plasmid # 54762 ; http://n2t.net/addgene:54762 ; RRID:Addgene_54762). The C-terminal degrons were appended to the eGFP gene by site-directed mutagenesis. Primers used for PCR (Suppl. Table 1) were designed to amplify an entire pBAD-eGFP plasmid while simultaneously adding degrons. The PCR products were then ligated and phosphorylated, and the obtained constructs were transformed into *E. coli* Top10 for selection on ampicillin.

Successfully transformed bacteria were used for plasmid isolation, and all constructs were verified by sequencing. The sequences of *clpP, clpX* and *sspB* were identical to the corresponding genes from strain K-12 substrain MG1655 which is widely used as a model *E. coli* strain (GenBank: U00096.3).

### Protein expression and purification

For *in vitro* studies protein expression was carried out in BL21 DE3 bacteria (for ClpX, ClpP, SspB), in *E. coli* Top10 strain (for eGFP-degrons) and in *ΔclpP* Keio collection strain (JW0427) (for ClpX, eGFP-degrons) using LB media with induction at OD_600_ ∼ 0.5 with 0.5 mM IPTG for pET28a constructs in BL21 DE3 strain or 0.02% arabinose for pBAD constructs in Top10 and *ΔclpP* strains. Overexpression of ClpP, SspB and eGFP-degrons was carried out at 30°C for 3h, ClpX at 18°C overnight, all at 140 rpm. Bacterial pellet was collected by centrifugation (5000rpm [5180xg], 30min, at 4°C) and frozen at -20°C for later use.

ClpX, ClpP, SspB and eGFP-degrons were purified on HisTrap HP 5 ml column (GE Healthcare) following the manufacturer’s guidelines. The lysis/loading buffer contained 50 mM Tris pH 8.0, 20 mM imidazole, 500 mM NaCl (for ClpX and eGFP-degrons) or alternatively 1 M NaCl (for ClpP and SspB) and with addition of 1 mM DTT for ClpX. Lysis by incubation for 30min (on a rotator at 4°C) with lysozyme (100 mg) and DNase (20U) was followed by sonication (15min, 45 on / 15 off sec, 40% amplitude at 4°C) and centrifugation (20000rpm [48380xg], 30 min, 4°C). Lysates were cleared by filtration (0.45μm) and applied on the HisTrap column. ClpX, ClpP and SspB elutions (using lysis/loading buffer supplemented to 1 M imidazole) were visualized on SDS-PAGE and proper fractions were selected, then combined and treated with SUMO protease for His-SUMO tag removal (2h on a rotator at 4°C). After cleavage, samples were concentrated in 10K MWCO centricons (PAL Corporation) (5000rpm [3214xg] at 4°C) and applied on a Superdex200 HiLoad 16/600 column (GE Healthcare) using 25 mM Hepes pH 7.6, 200 mM KCl, 5 mM MgCl_2_, 10% glycerol buffer with the addition of 0.5 mM of TCEP in case of ClpX. The eGFP-degrons were purified on HisTrap and Superdex200 columns in a tandem configuration without purification tag (His-TEV) removal in between. Based on SDS-PAGE, selected SEC fractions were combined, and protein concentration was measured using the Bradford method (for ClpX) or based on the UV absorbance profile (for ClpP, SspB, His-TEV-eGFP-degrons, further denoted as eGFP-degrons). All proteins were flash frozen in liquid nitrogen and stored at -80°C for later experiments.

### ATP activity assay

*In vitro* ATP hydrolysis by ClpX was measured using an NADH-coupled assay as previously described (Nørby, 1988). A mixture of ClpX interacting protein (0-27.4 μM SspB_2_, eGFP-SsrA or eGFP-SsrA with constant concentration of 2.5 μM SspB_2_), 2.5 mM ATP, 2.5 mM phosphoenolpyruvate, 50 µg/ml pyruvate kinase, 1 mM NADH, and 50 µg/ml lactate dehydrogenase in the reaction buffer (50 mM Tris-HCl pH 7.5, 200 mM NaCl, 5 mM MgCl_2_) was incubated at 30°C for 30 min. As a negative control for the study of the eGFP-ssrA interaction, an eGFP protein not fused to ssrA degron was used. The assay was started by the addition of ClpX_6_ to a final concentration of 0.08 µM in the 100 µl final reaction volume. Changes in NADH concentration were monitored by measuring absorbance at 340 nm for 1.5h (in 1 min intervals) at 30°C in a Tecan Infinite 200 Pro plate reader instrument. All measurements were performed at least in a triplicate. The rate of ADP formation was calculated from the linear loss of fluorescence assuming a 1:1 correspondence between ATP regeneration and NADH oxidation and a Δε_340_ of 6.23 µM^-1^cm^-1^. Curve fitting and statistical analysis (one-sample and unpaired two-sample two-tailed t-test) was performed via Graphpad Prism 9 software (Graphpad Software, LLC).

### Native electrophoresis

*In vitro* protein-protein interaction studies were performed by means of non-denaturing polyacrylamide gel electrophoresis. Individual proteins (8 μM for eGFP-degron and 8 μM for SspB_2_) or protein mixtures (8 μM each) were incubated prior to electrophoresis for approximately 10 min at RT in the reaction buffer (50 mM Tris-HCl pH 7.5, 200 mM NaCl, 5 mM MgCl_2_). Protein samples were separated on 15% native polyacrylamide gels in Tris/borate buffer and the results visualized by staining the gels with SimplyBlue Safe Stain (Invitrogen, LC6060).

### Microscale Thermophoresis (MST)

For measurements of binding between SspB and eGFP-degrons, 100 nM eGFP-degrons and a series of 16 two-fold dilutions of SspB ranging from 7.45 nM to 244 μM in MST buffer (25 mM Hepes 7.6, 200 mM KCl, 5 mM MgCl_2_, 10 % glycerol) supplemented with 0.05 % Tween 20 were prepared. Then each ligand dilution was mixed with one volume of eGFP-degron, which led to the final eGFP-degron concentration of 50 nM and final SspB concentrations ranging from 3.72 nM to 122 μM. After 15 min incubation at RT, the samples were loaded into Monolith NT.115 Premium Capillaries (NanoTemper Technologies), and measurements were carried out using the Monolith NT.115 instrument (NanoTemper Technologies) at 20°C. Instrument parameters were adjusted to 20 % LED power and medium MST power.

All measurements were performed at least in triplicates. Protein binding parameters were calculated using the signal from an MST-on time of 10 s. Curve fitting and Scatchard plots, and statistical analysis of the obtained constants (one-sample two-tailed t-test) were performed via Graphpad Prism 9 software (Graphpad Software, LLC).

### Fluorescence-based measurements of *in vitro* degradation

*In vitro* degradation of eGFP-degron by ClpXP analysis was performed in the 100 µl final volume in the reaction buffer (50 mM Tris-HCl pH 7.5, 200 mM NaCl, 5 mM MgCl_2_). A mixture of 0.08 µM ClpX_6_, eGFP-degron (0-10 μM), SspB (2.5 μM SspB_2_, if added), 4 mM ATP, 2.5 mM creatine phosphate, and 50 µg/ml creatine kinase was incubated at 30°C for 30 min. As a negative control, the eGFP protein not fused with any degron was used. The degradation assay was started by the addition of 0.21 µM ClpP_14_ for the final protease machinery assembly in ClpX_6_:ClpP_14_ = 3:8 ratio. Changes in eGFP fluorescence (at excitation λ=489 nm and emission λ=550 nm) were monitored for 1.5 h at 30°C in a Tecan Infinite 200 Pro plate reader instrument. All measurements were performed at least in triplicates. Degradation rates were calculated from the initial linear loss of eGFP fluorescence. Curve fitting using the Michaelis-Menten equation and statistical analysis (one-sample and unpaired two-sample two-tailed t-test) was performed via Graphpad Prism 9 software (Graphpad Software, LLC).

### SDS-PAGE analysis of *in vitro* degradation

Reactions for SDS-PAGE analysis of *in vitro* eGFP-degron degradation by ClpXP were prepared the same way as for fluorescence readouts, with eGFP-degron or eGFP concentration set at 1 μM. 10 µl samples were collected in 15 min intervals up to 120 min (with additional time points for AANDENYSENYALAA of 3, 6, 9 and 12 min). The reaction was stopped by adding the SDS-PAGE loading buffer followed by 5 min incubation at 95°C. Prepared samples were applied on 12% SDS-PAGE gels, which were developed at 200 V. Resolved proteins were visualized using the SafeStain method.

### Time-course measurements of *in vivo* degradation

Precultures were prepared by inoculating a small amount of a glycerol stock of *E. coli* BW25113 (previously transformed with pBAD-eGFP constructs) in liquid LB medium with ampicillin (50 μg/ml) and incubating overnight at 37°C in an orbital shaker. The following day, the precultures were diluted 50 times in fresh medium and the cultures were grown to mid-exponential phase. The expression of eGFP-degron fusions was induced by addition of L-arabinose to the final concentration of 0.001%. The induced cultures were incubated overnight in 18°C. Bacteria were then diluted 10 times by adding 20 μl of the cultures to 180 μl of M9 medium supplemented with glucose (5%), thiamine (10 μg/ml), biotin (10 μg/ml), trace elements, and additionally spectinomycin (100 μg/ml) to stop protein translation. The measurements of OD_600_ and fluorescence (excitation at λ = 489 nm and emission at λ = 520 nm) were made every 15 minutes for 6 hours using Tecan M200 Pro plate reader at 30°C. The fluorescence results were normalized to the optical density. The measurements were further normalized by setting the values at zero time-points as 100%.

### Plate spot assay of *in vivo* degradation

2-3µL drops of 100x diluted overnight cultures (prepared as described above) were placed in triplicates or quadruplicates on LB agar plates supplemented with 0.001% L-arabinose and antibiotics: ampicillin (50 μg/ml) for WT strain (BW25113), and ampicillin (50 μg/ml) with kanamycin (15 μg/ml) for the deletant strains (*ΔclpX* (JW0428-KC), *ΔclpP* (JW0427-KC) and *ΔsspB* (JW0866-KC)). The plates were incubated overnight at 37°C. Images of the plates were taken with the Bio-Rad Chemidoc Imager using a fluorescein filter and analyzed with Fiji software.

## RESULTS

### Optimization of protein preparation for ClpX, ClpP, SspB and eGFP-degron constructs

To be able to compare the efficiency of studied degrons in protein degradation *in vitro*, it was crucial to obtain a fully functional ClpX-ClpP-SspB complex. The first step to achieve this goal was to produce full-length recombinant proteins, especially ClpX together with the Zinc Binding Domain (ZBD), which is necessary for the interaction of ClpX with SspB.

In order to obtain the highest possible consistency of results, all proteins of the ClpX-ClpP-SspB complex were cloned in our lab from Top10 *E. coli* strain (*Escherichia coli*; K-12 substr. MG1655) into a pET28a vector containing N-terminal His-SUMO tags. After screening for the best overexpression conditions in *E. coli* BL21 DE3 strain, ClpP and SspB proteins were deemed sufficiently expressed after a few hours at 30°C, whereas ClpX as the most temperature-sensitive protein was expressed overnight at 18°C. Substrate proteins comprising eGFP fused with a C-terminal ssrA-derived degron (“eGFP-degrons”) and N-terminal His-TEV purification tags were expressed from pBAD vectors to enable their controlled induction in *E. coli* BW25113 strain and its derivatives.

ClpX, ClpP and SspB were purified by affinity chromatography (HisTrap column) followed by His-SUMO tag removal and size exclusion chromatography (Superdex200 column). eGFP-degrons were purified using the same columns in a tandem setup, without any His-TEV tag removal step.

All proteins were purified with good efficiency, although, even after several attempts to optimize the protocols, significant degradation for ClpX and eGFP-degrons was still present (Supplementary Fig. 1). To overcome this problem, His-SUMO-ClpX fusion was recloned into the pBAD vector enabling expression in the *ΔclpP E. coli* strain that has reduced protein degradation due to the lack of ClpP peptidase. For the same reason, eGFP-degron expression was also finally switched to the Δ*clpP* strain. These changes in expression protocols resulted in good protein expression levels and a significant reduction in the observed degradation (Fig. 1C).

### Validation of ZBD and ClpX active site function by an ATPase activity assay

To determine if the purified ClpX protein is active *in vitro*, ATP hydrolysis by ClpX was measured using an NADH-coupled assay (Fig. 2). Several experimental conditions were evaluated: with increasing concentration of SspB or ssrA, and alternatively with an increasing concentration of eGFP-ssrA at the constant concentration of 2.5 µM SspB_2_. ClpX was active and SspB alone increased reaction velocities up to roughly two-fold (K = 0.22 ± 0.03 µM with the half-maximal value at 3.13 µM). The presence of the eGFP-ssrA substrate increased reaction velocities even further at larger eGFP-ssrA concentrations (K = 0.50 ± 0.01 µM with the half-maximal value of 1.38 µM). Additionally, an expected increase in both the half-maximal value and reaction velocities was observed in the presence of both SspB and eGFP-ssrA (K = 0.20 ± 0.03 µM with the half-maximal value of 3.42 µM), indicating the correct interaction of SspB with the ZBD domain of ClpX.

**Figure 2.**
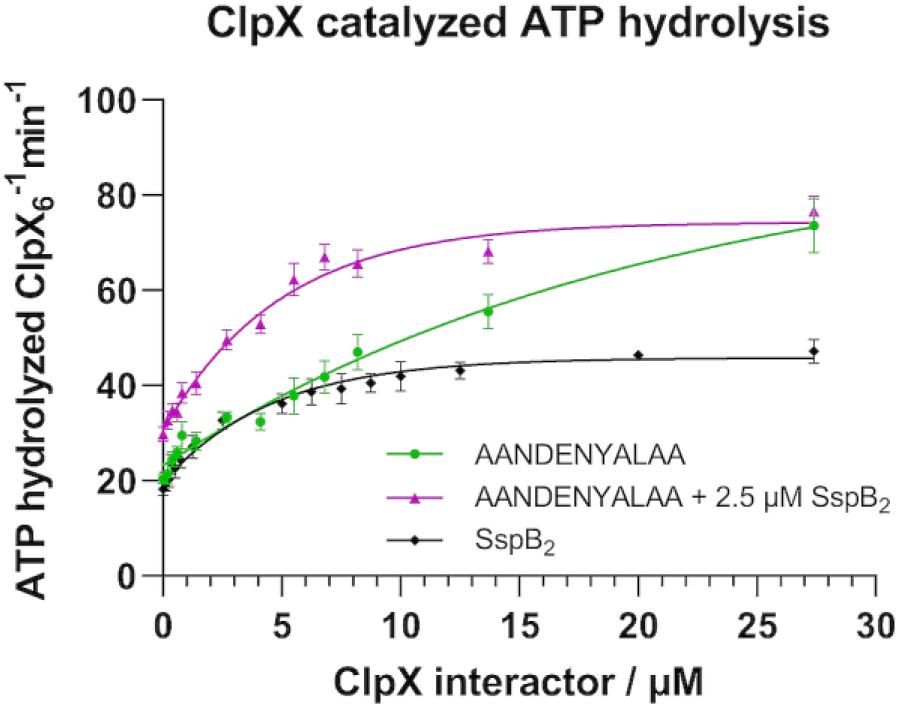
The *in vitro* assay of ClpX ATPase activity. ATP hydrolysis catalyzed by ClpX was measured using the NADH coupled assay by monitoring changes in NADH absorbance as a function of time in the presence of increasing concentrations of SspB_2_ (black curve) and increasing concentrations of eGFP-AANDENYALAA alone (green curve) or with the constant addition of 2.5 µM SspB_2_ (violet line). Data are presented as means with error bars as SEM for at least three technical replicates.

### The comparison of binding of eGFP-ssrA variants by SspB

The next step was to check whether the purified SspB protein interacted *in vitro* with the individual ssrA variants. For this purpose, native electrophoresis was used to visualize the interaction of various eGFP-degrons with SspB_2_. Two groups of degrons emerged in this comparison (Fig. 3A, B). The first group consisted of degrons that bound SspB due to the presence of the AANDENY sequence responsible for the interaction of the ssrA tag with SspB, and the second group comprised degrons that didn’t bind SspB and were correspondingly lacking the AANDENY sequence. To quantify the differences in the binding efficiency of individual degrons to SspB, we determined their binding constants using microscale thermophoresis (MST) (Fig. 3C, D and Supplementary Fig. 2.). MST confirmed not only that the AANDENY sequence is essential for degrons to bind to SspB, but also demonstrated that all degrons containing the AANDENY sequence bind with comparable efficacy to WT ssrA (K_D_ = 0.08 ± 0.02 µM) ranging from K_D_ = 0.10 ± 0.02 µM for AANDENYSENY to K_D_ = 0.33 ± 0.06 µM for AANDENYADAS. The two degrons with the mutated “ADAS” C-terminal sequence replacing ALAA (AANDENYADAS; AANDENYSENYADAS: K_D_ = 0.30 ± 0.08 µM) showed 2-3 fold higher dissociation constants, but within the same order of magnitude as the other AANDENY-containing sequences. The binding of degrons lacking an AANDENY sequence was not detected under our experimental conditions.

**Figure 3.**
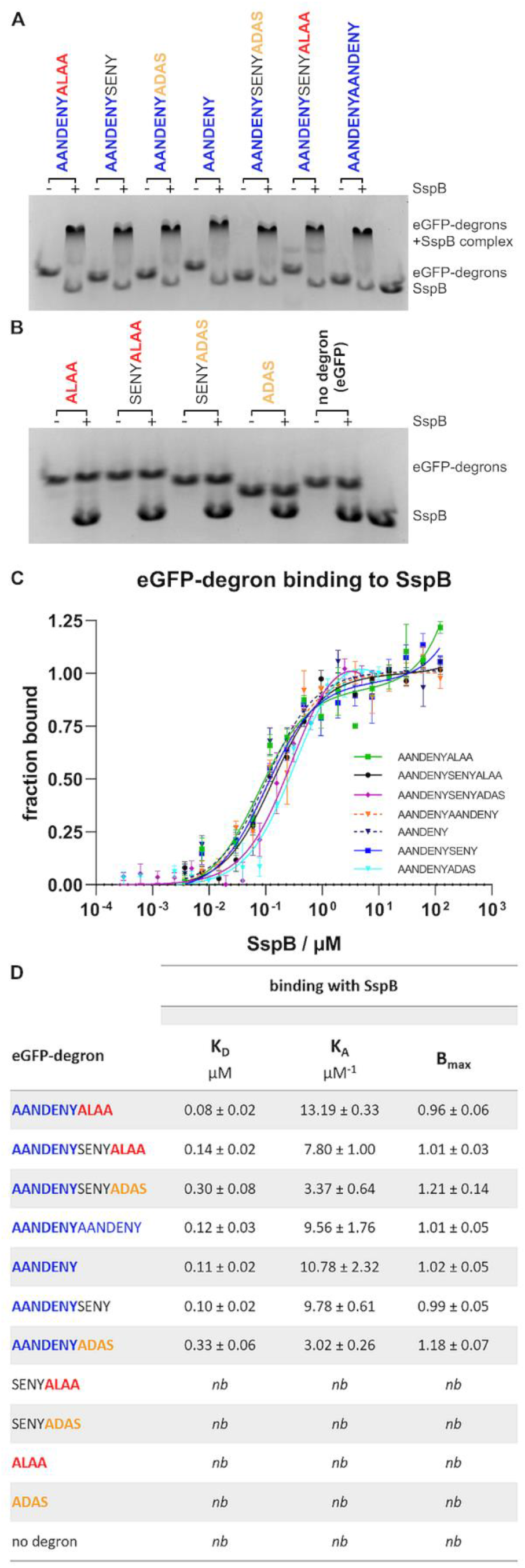
The *in vitro* study of SspB binding to eGFP-degron proteins. (A, B) Native electrophoresis revealed formation of complexes between SspB and eGFP proteins coupled with degrons containing the AANDENY motif (A), in contrast to no visible binding of SspB to eGFP proteins coupled with degrons lacking the AANDENY motif (B). (C) Confirmation of SspB binding to eGFP with degrons containing the AANDENY motif by MST.). Data are presented as means with error bars as SEM for at least three technical replicates. (D) Quantification of the binding parameters (K_D_, K_A_, and B_max_) based on the analysis of MST data. nb – no detectable binding. Data are presented as mean ± SEM.

### *In vitro* degradation of eGFP-degrons by the ClpXP complex

Degradation of eGFP-degrons by the ClpXP complex was measured as a decrease of eGFP fluorescence using a plate reader and applying a 3:8 ratio of ClpX_6_:ClpP_14_ (Fig. 4). In the absence of SspB, K_M_ could be determined only for three degrons (AANDENYALAA: K_M_ = 3.38 ± 0.39 µM; AANDENYSENYALAA: K_M_ = 1.55 ± 0.20µM; SENYALAA: K_M_ = 4.31 ± 0.69 µM), confirming that the ALAA sequence was essential for degradation and the longer the sequence preceding the ALAA fragment, the more efficient the degradation was. In the presence of SspB, K_M_ could be evaluated for 6 degrons and showed notable improvement in each case, up to by an order of magnitude for the WT ssrA sequence. The most efficient degradation was observed for degrons containing both the AANDENY and ALAA sequences (AANDENYALAA: K_M_ = 0.47 ± 0.06 µM; AANDENYSENYALAA: K_M_ = 0.48 ± 0.05 µM), with no effect of the additional linker (SENY). The degron with the mutated “ADAS” C-terminal sequence replacing ALAA (AANDENYSENYADAS: K_M_ = 0.79 ± 0.16 µM) was less effective, especially for the sequence without the SENY linker (AANDENYDAS: K_M_ = 7.8 ± 2.9 µM). Degrons containing the AANDENY sequence needed a longer C-terminal extension (AANDENYAANDENY: K_M_ = 1.41 ± 0.28 µM; AANDENYADAS: K_M_ = 7.8 ± 2.9 µM) to work efficiently. It was impossible to determine K_M_ for the degron consisting only of the ALAA sequence unless it had a preceding linker (SENYALAA: K_M_ = 2.61 ± 0.39 µM). For the rest of the degrons, K_M_ could not be determined due to a lack of sufficient degradation.

**Figure 4.**
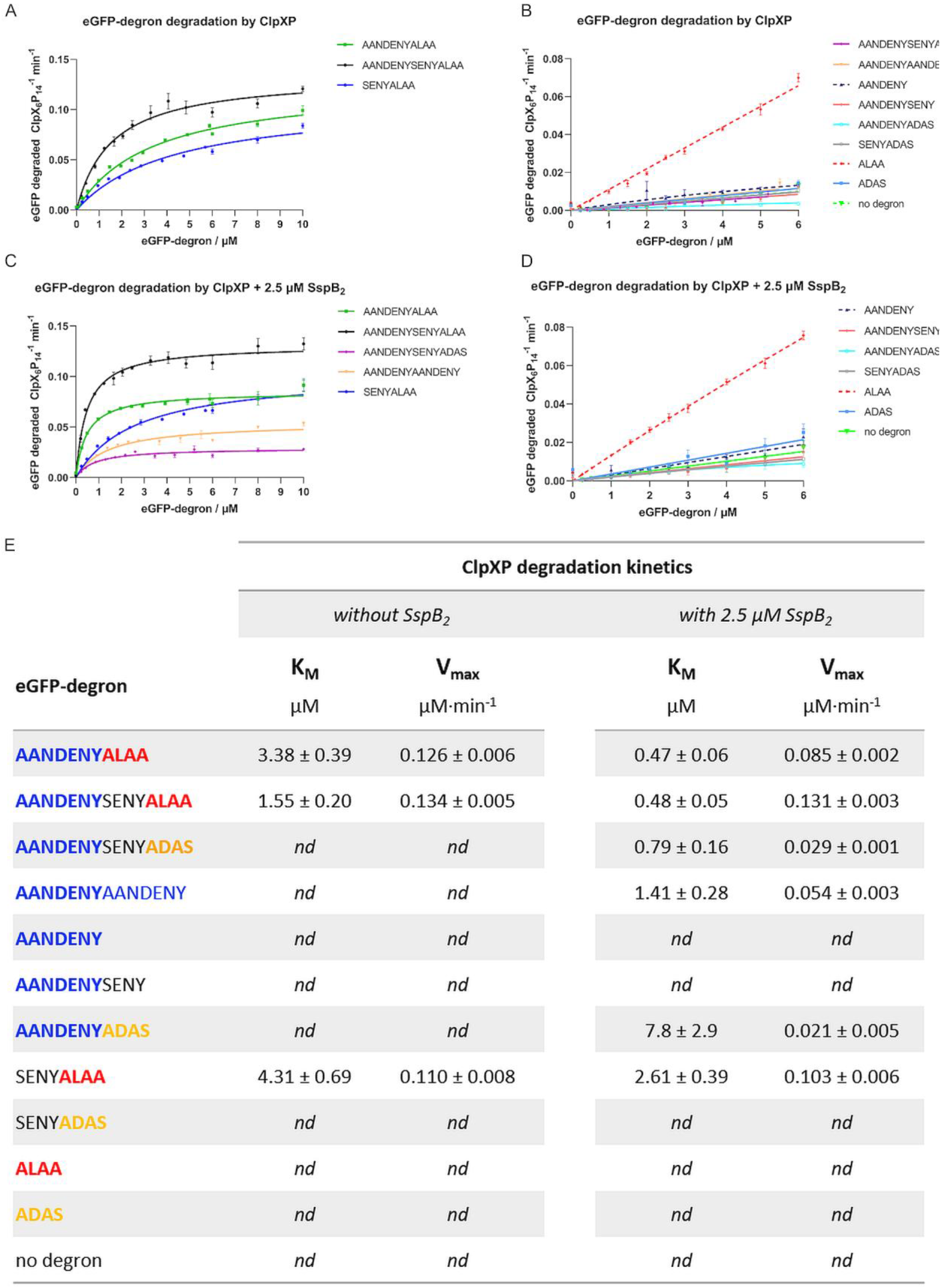
*In vitro* degradation of eGFP-degron proteins by ClpXP. Evaluation through changes in the eGFP fluorescence in the absence (A, B) or presence of 2.5 µM SspB_2_ protein (C, D). Data are presented as means with error bars as SEM for at least three technical replicates. (E) The initial rate of protein degradation per ClpX_6_P_14_ unit at different eGFP-degrons concentration was fit to the Michaelis-Menten plot allowing the determination of the process kinetic parameters (K_M_, V_max_) in the absence (left) or presence of 2.5 µM SspB_2_ (right); nd – no discernible degradation. Data are presented as mean ± SEM for at least three technical replicates.

To confirm the stability of the ClpX-ClpP-SspB *in vitro* degradation system itself, degradation was additionally analyzed using SDS-PAGE for the most efficient AANDENYALAA and AANDENYSENYALAA degrons. Substrate removal was more efficient for AANDENYSENYALAA (∼15 min to reach the steady-state level) in comparison with AANDENYALAA (∼60 min to reach the steady-state level) (Fig.5 A, B). A small, residual amount of substrates present until the end of the reaction likely resulted from the prior spontaneous degradation of eGFP-degrons to the eGFP itself, which is not degraded further. No additional degradation of SspB, ClpXP or the regeneration system components was observed after the degradation of eGFP-degrons, confirming specific substrate removal and stability of the applied assay.

**Figure 5.**
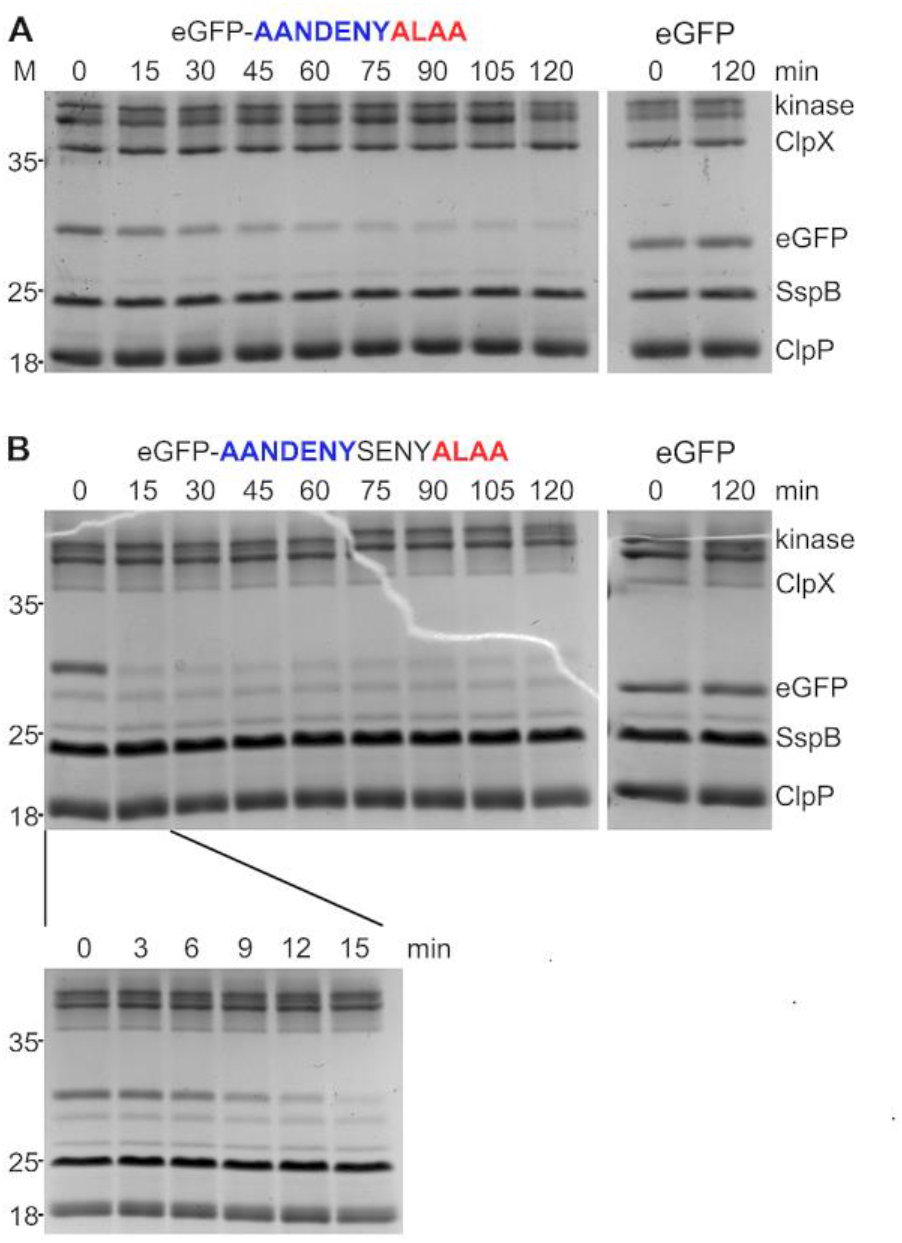
SDS-PAGE analysis of the *in vitro* eGFP-degron degradation by ClpXP. The progress of eGFP-AANDENYALAA (A) and eGFP-AANDENYSENYALAA (B) degradation in the presence of 2.5 µM SspB_2_ protein was sampled over a time course of 2h. Degradation of eGFP without a degron tag served as a control (right).

### *In vivo* degradation of eGFP-degrons

Our results of the *in vitro* degradation assays obtained for eGFP-degrons were confirmed by additional *in vivo* analyses carried out in bacterial cultures. Two methods were used: an analysis of changes in eGFP-degron fluorescence either in liquid culture over time (Fig. 6A) or in a spot assay on agar plates (Fig. 6B). The most efficient method of comparing degron efficiency was an estimation of the level of eGFP fluorescence of bacterial colonies overexpressing eGFP-degrons in the plate spot assay (Fig. 6B). The fluorescence level of eGFP-degrons was determined for WT strain and three deletant strains *ΔclpX, ΔclpP* and *ΔsspB* to establish whether ClpX, ClpP or SspB would affect degradation. In the WT strain, eGFP-degron degradation resulted in the fastest and largest decrease in fluorescence for AANDENYALAA, ALAA, SENYALAA, and AANDENYSENYALAA degrons (Fig. 6A, C). A slower decrease was observed for AANDENYAANDENY, AANDENYSENYADAS, and for the more lagging AANDENYADAS (Fig. 6A), correlating also with the lower fluorescence level in the spot assay for these three degrons (Fig. 6B). AANDENY, AANDENYSENY, ADAS, and SENYADAS degrons resulted in only slight decrease in fluorescence, with AANDENY matching the most closely the levels of eGFP without any degron tag (Fig. 6A, C). The overall trends in fluorescence levels were similar in the WT, *ΔclpX* and *ΔsspB* strains (Fig. 6C, D, F), with ALAA-containing degrons causing the most efficient eGFP degradation, and this reduction was the most evident in the *ΔsspB* strain. For the *ΔclpP* strain, substantial degradation was observed only for the eGFP-ALAA fusion (Fig. 6B, E).

**Figure 6.**
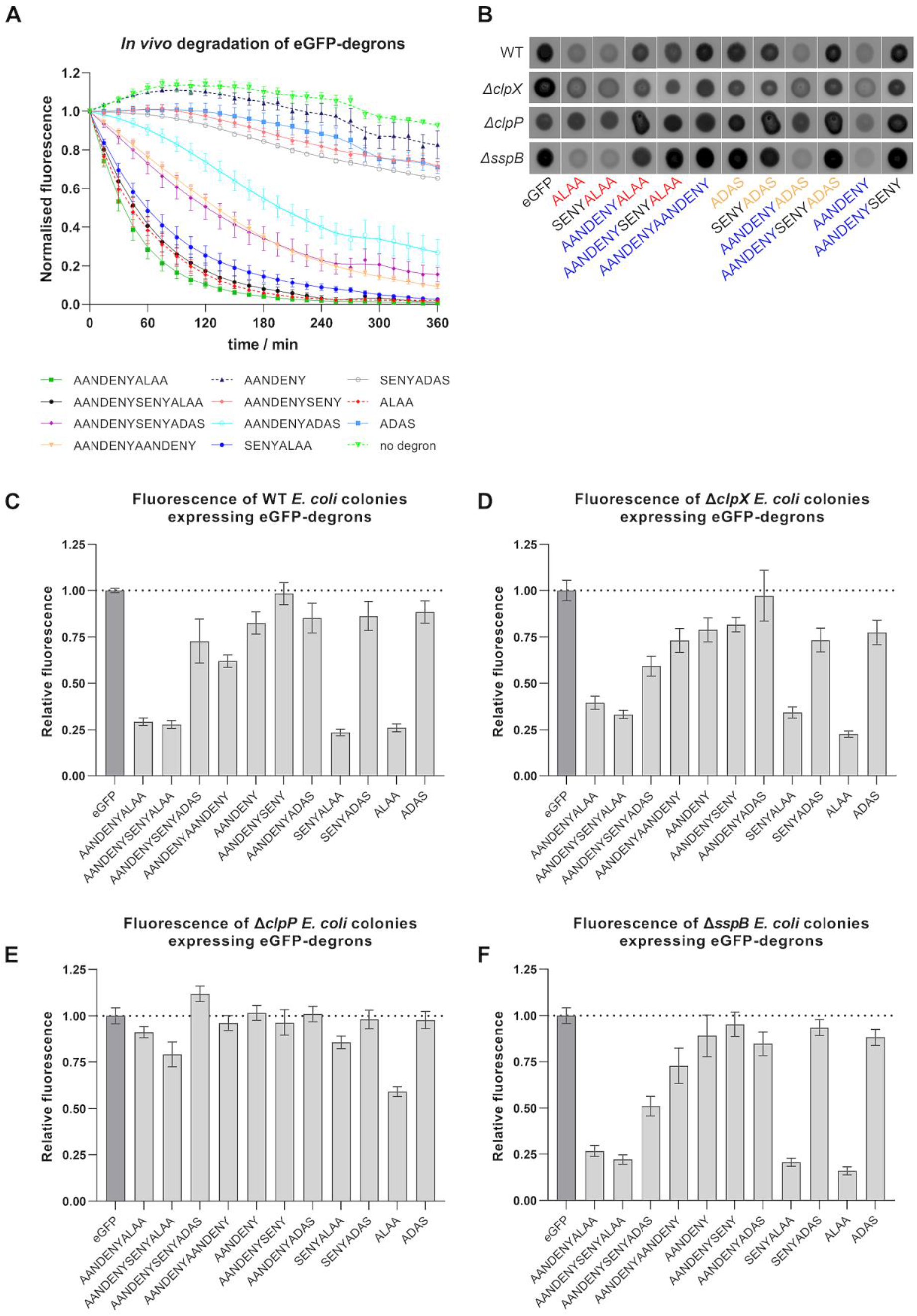
*In vivo* degradation of eGFP-degrons by ClpXP. (A) Changes in fluorescence observed for WT strain overexpressing eGFP-degrons over a time course of 6h. (B) A representative set of images of the bacterial colonies in the plate spot assay for the fluorescence phenotype (eGFP fluorescence signal shown in black). The fluorescence signal intensity was quantified, normalized to eGFP control and presented as relative fluorescence for the WT (C), *ΔclpX* (D), *ΔclpP* (E) and *ΔsspB* (F) strains. Data are presented as means with error bars as SEM for at least three biological replicates.

## DISCUSSION

For the purpose of comparing and characterizing the properties of ssrA variants, it was necessary to set up expression and purification protocols for all eGFP-degrons and the components of the ClpX-ClpP-SspB complex, as well as methods required to check their activity and interactions. The expression of proteins at the beginning in the Top10 for eGFP-degrons and in the BL21 DE3 strain for ClpX, ClpP and SspB unfortunately resulted in a noticeable degradation in the case of eGFP-degrons and ClpX. In the case of ClpX, which in our hands was generally the most challenging protein to purify, it could be attributed to the high mobility of the ZBD domain, which can be easily digested by various proteases. Similar problem was encountered previously in the structural studies of ClpX. Typically, ClpX structures were obtained without the ZBD domain (Fei *et al*, 2020), and if ZBD was present, the resolution for it was much lower than for the rest of the ClpX structure (Gatsogiannis *et al*, 2019). For eGFP-degrons the observed degradation likely took place during their expression in bacteria, proving efficient *in vivo* degradation of proteins with ssrA tag or its modifications. Only switching to expression in the *ΔclpP* strain allowed to obtain proteins that did not show significant degradation.

In the next step, we ascertained that the obtained ClpX was active in an ATPase assay. Our results confirmed the previously reported (Wah *et al*, 2002; Song & Eck, 2003) positive effect of the SspB adaptor on the ATPase activity of ClpX. Since ZBD is explicitly necessary for the interaction with SspB (Park *et al*, 2007), this strongly indicated the presence of a functional ZBD domain within the ClpX protein, with the downstream consequences on degradation by improving substrate engagement by ClpX.

The analysis of binding of eGFP-degrons to SspB protein studied on native gels confirmed the previously reported results of a mutational analysis of the GFP-ssrA tag (Flynn *et al*, 2001) that AANDENY sequence is necessary for eGFP-degrons to bind SspB. Our MST measurements allowed for a more detailed binding analysis and the determination of the K_D_ for eGFP-ssrA and SspB at ∼80 nM. Similar K_D_ values were observed previously in ITC measurements for GFP-ssrA and SspB, where the binding constant was determined to be 75 ± 30nM (Hersch *et al*, 2004) or even 16 ± 4 nM (Wah *et al*, 2002). The application of the eGFP-degron fusions allowed us not only to evaluate the interactions using MST but also to reduce the K_D_ value by one order of magnitude compared to the isolated ssrA tag, which is known to bind more weakly (Wah *et al*, 2002). Additionally, MST as a relatively high - throughput and precise method allowed us to determine even minor differences in the binding of the SspB protein to eGFP-degrons. We noted that the ADAS sequence caused a slight decrease in the eGFP-degron binding by SspB in contrast to the wt ALAA end, despite the expected independence of the AANDENY and ALAA modules in binding to SspB and ClpX, respectively. This difference may be due to steric requirements or electrostatic interactions related to the presence of the negatively charged aspartate in the DAS fragment. To our knowledge, this is the first time that the DAS fragment has been reported to negatively influence SspB binding since, typically, the C-terminal amino acids of ssrA have been tested mostly for recognition by ClpX (McGinness *et al*, 2006). It is not clear whether the disruptive effect of DAS would have any physiological relevance since it was observed under isolated conditions; however, it could be of importance in the design of AANDENY-containing modules used to mediate interactions with the SspB protein. Our results suggest that the sequence that follows the AANDENY fragment should also meet specific requirements for the most efficient binding of the substrate protein by SspB. The overall degron length in terms of the extensions beyond the AANDENY motif did not matter in SspB binding, even in the case of the eGFP-AANDENYAANDENY variant which seemed to bind better to SspB but without a significant change of the K_D_. This may suggest the impossibility of occupying more than one binding site by SspB on such a short peptide regardless of the presence of two SspB binding sites. This result may also indicate relatively slow dissociation of the complex between SspB and the AANDENY motif, which would render the second binding motif present in AANDENYAANDENY less relevant for improving SspB binding despite the increased local concentration of binding sites.

The key part of our work was comparing the degradation effectiveness of eGFP-degrons by the ClpXP complex and the effect of SspB on it. The application of our standardized reaction conditions throughout our assays to a whole set of ssrA tag variants made it easier to compare their K_M_ values and draw conclusions which parts of degrons are necessary for degradation, which can be replaced, and which are largely negligible. In the absence of SspB, degradation was observed only for eGFP-degrons containing the ALAA end preceded with an extending fragment such as the SENY linker. The longer the preceding fragment, the more efficient the degradation was observed. The ALAA fragment alone was insufficient for degradation. The ADAS fragment which is known to interact weaker with ClpX (McGinness *et al*, 2006), rendered eGFP fusions retardant to degradation regardless of the extension preceding ADAS, even for the longest AANDENYSENYADAS degron variant.

In the presence of SspB, the K_M_ values for eGFP-degrons, which were partially degraded without SspB, were further lowered. Additionally, degradation was induced for 3 more eGFP-degrons (AANDENYADAS, AANDENYSENYADAS, AANDENYAANDENY), proving the positive influence of the AANDENY motif on the degradation efficiency and even compensating for the weaker binding of the mutated ADAS equivalent of ALAA. Interestingly, eGFP-AANDENYAANDENY was degraded efficiently despite the lack of the ClpX binding motif (ALAA or ADAS), suggesting that the terminal repeat of the AANDENY fragment provided sufficient reach to bind nonspecifically to ClpX for degradation. The decrease in K_M_ was also observed for the eGFP-SENYALAA, which does not have the AANDENY motif and did not bind to SspB in our hands. This improvement in degradation efficiency can be attributed to the increased ClpX activity in the presence of SspB, as observed in our ATPase activity assays. Such a boost could result in the increased degradation of eGFP-SENYALAA by ClpX without direct SspB binding to the degron. However, the increased ATPase activity was not sufficient to efficiently degrade short degrons or degrons lacking the ALAA fragment. In summary, the eGFP-degron must be able to engage ClpX and be at least slightly degradable in the absence of SspB so that the increased ATPase activity of ClpX in the presence of SspB could positively influence its degradation. The positive effect of SspB observed by us on the degradation efficiency is consistent with the literature data (Wah *et al*, 2002; McGinness *et al*, 2006, 2007). The slight differences in the intensity of its effect may be attributed to the differences in the experimental conditions, various concentrations of proteins used in the reaction mixture and the various ClpX/ClpP/SspB ratios. Despite these differences, our results and the literature data (McGinness *et al*, 2007) similarly indicate that the highest impact of SspB protein on the eGFP-degron degradation efficiency was observed for the eGFP-AANDENYSENYADAS variant.

The results obtained *in vivo* were consistent with our *in vitro* results showing that eGFP-degrons containing the ALAA sequence were degraded the most efficiently. Significantly reduced fluorescence was observed for those variants (AANDENYALAA, AANDENYSENYALAA and SENYALAA), for which eGFP-degrons were also degraded *in vitro* even in the absence of SspB. For the remaining eGFP-degrons, the decrease of fluorescence was not pronounced, confirming again *in vitro* degradation requirements for the proper degron length and the ALAA fragment. An additional common element for our *in vitro* and *in vivo* results was the atypical behaviour of the eGFP-ALAA fusion, which *in vitro* was degraded with an intermediate efficiency among our set of degrons even in the absence of SspB, but with a linear dependence on the tested substrate concentration that prevented K_M_ calculations. *In vivo* the eGFP-ALAA was degraded very efficiently independently of all the factors we addressed. This was not due to the low level of eGFP-ALAA expression which was deemed sufficiently high by our time-course experiments, but rather that it could be degraded as well by other proteases outside the ClpXP pathway. This could result from a cross-talk of the ssrA system with other pathways in which hydrophobic or alanine-rich degrons are recognized as signals for degradation. The involvement of other proteases is likely if we take into account that in the *ΔclpX* strain some eGFP-degrons were still degraded, whereas in the *ΔclpP* strain only the aforementioned eGFP-ALAA was depleted. Our results indicate that in BW25113 cells, ClpX is redundant for ssrA-tagged substrates and could be successfully replaced by another unfoldase such as ClpA, while the absence of ClpP unequivocally limits degradation of almost all of the studied degrons. Degradation observed in the *ΔsspB* strain was similar to that observed for WT and *ΔclpX* strains, corroborating the redundancy of the ClpX pathway or the SspB itself. Recently, it was shown that the *E. coli* ribosome-associated trigger factor (TF), a chaperonin which is highly abundant in the cell and acts early during the folding process, enhances degradation *in vitro* and *in vivo* of various ClpXP complex substrates including ssrA-taged proteins (Rizzolo *et al*, 2021). The role of TF as an additional adaptor could contribute to the lack of an effect of *sspB* deletion on our tested eGFP-degrons.

The results and protocols presented here can serve as a benchmark for designing new degrons as well as providing guidelines on their comparison. However, the degron research strategies we used also have their limitations. Firstly, the complicated stoichiometry of oligomeric components may pose a problem. Changes in stoichiometry can cause changes in degradation kinetics (especially if the substrate is also an oligomer) or in the selectivity of the degradation system. Secondly, it is crucially important to test degron performance in the case of individual substrates/targeted proteins, as the substrate itself may alter degron availability and thus affect the efficiency of degron binding or its degradation. With these difficulties in mind, we aimed at establishing the most robust and straightforward strategies and methods, which allowed us to discover subtle differences between degrons in terms of their binding strength or degradation efficiency.

Building on this type of knowledge, degron variants can be fine-tuned for two basic types of applications. We ranked the propensity of the tested ssrA variants for SspB binding and for eGFP-degron degradation (Fig. 7), which should facilitate the choice of the degron appropriate to the desired task, such as efficient degradation of the fusion protein or only its promoted interaction with SspB. The best binding to SspB is provided by the wt ssrA sequence, but if more efficient degradation is needed, especially in the absence of SspB, then AANDENYSENYALAA would be a better choice. AANDENY or AANDENYSENY degrons would be the most suitable choice for efficient tag binding without degradation. Our results confirm that AANDENYSENYADAS and AANDENYADAS enable degradation which might be regulated by SspB presence (Griffith & Grossman, 2008; Davis *et al*, 2011; Kim *et al*, 2011).

**Figure 7.**
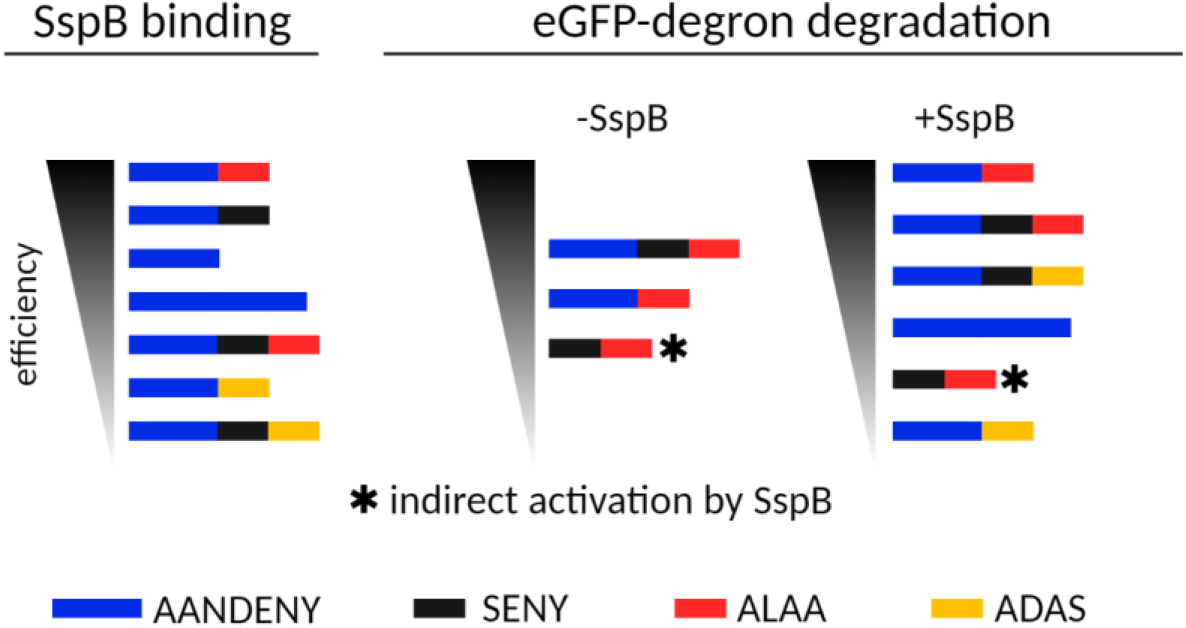
A schematic representation of eGFP-degron properties as ranked by their SspB binding efficiency and eGFP-degron degradation.

The ssrA degron is often used simply as an interaction module, for example in photoswitches (Guntas *et al*, 2015). Degrons can also be exploited as protease recruiting ligands in the design of degrader molecules in the targeted protein degradation field. For the latter purposes, it is essential to distinguish the effects of degron sequence and length on adaptor and protease binding from those parameters that lead to degradation and depletion of the degron-based molecular tools. Our benchmark of methods and protocols may serve the budding targeted protein degradation studies in bacteria in establishing more stable and efficient bacterial degraders.

## Acknowledgements

We would like to thank Rafał Augustyniak for helpful discussions. This work was financed under the grant agreement POIR.04.04.00-00-5EC1/18-00 for the project “Proteolysis-targeting strategies in bacterial systems for functional studies of proteins and improvement of antibiotics” carried out within the FIRST TEAM programme of the Foundation for Polish Science co-financed by the European Union under the European Regional Development Fund. Work in the laboratory was supported by the EMBO Installation grant to MWG. MWG is the recipient of the L’Oréal-UNESCO For Women in Science scholarship from L’Oréal Poland and the Ministry of Education and Science, Poland.

## Author Contributions

MMK purified proteins and performed in vitro degradation assays. AA performed ATPase, binding and in vitro degradation assays and statistical analysis. MAI performed cloning and in vivo assays and prepared vector graphics. PES performed in vivo assays. PKT and CD performed MST measurements. MWG acquired funding and supervised the project team. MMK wrote the first draft of the manuscript. MAI, AA, MWG wrote sections of the manuscript. AA and MAI contributed equally to the manuscript. All authors contributed to the conception and preparation of the manuscript and approved the submitted version.

## SUPPLEMENTARY MATERIAL

### SUPPLEMENTARY TABLES

**Supplementary Table 1.**
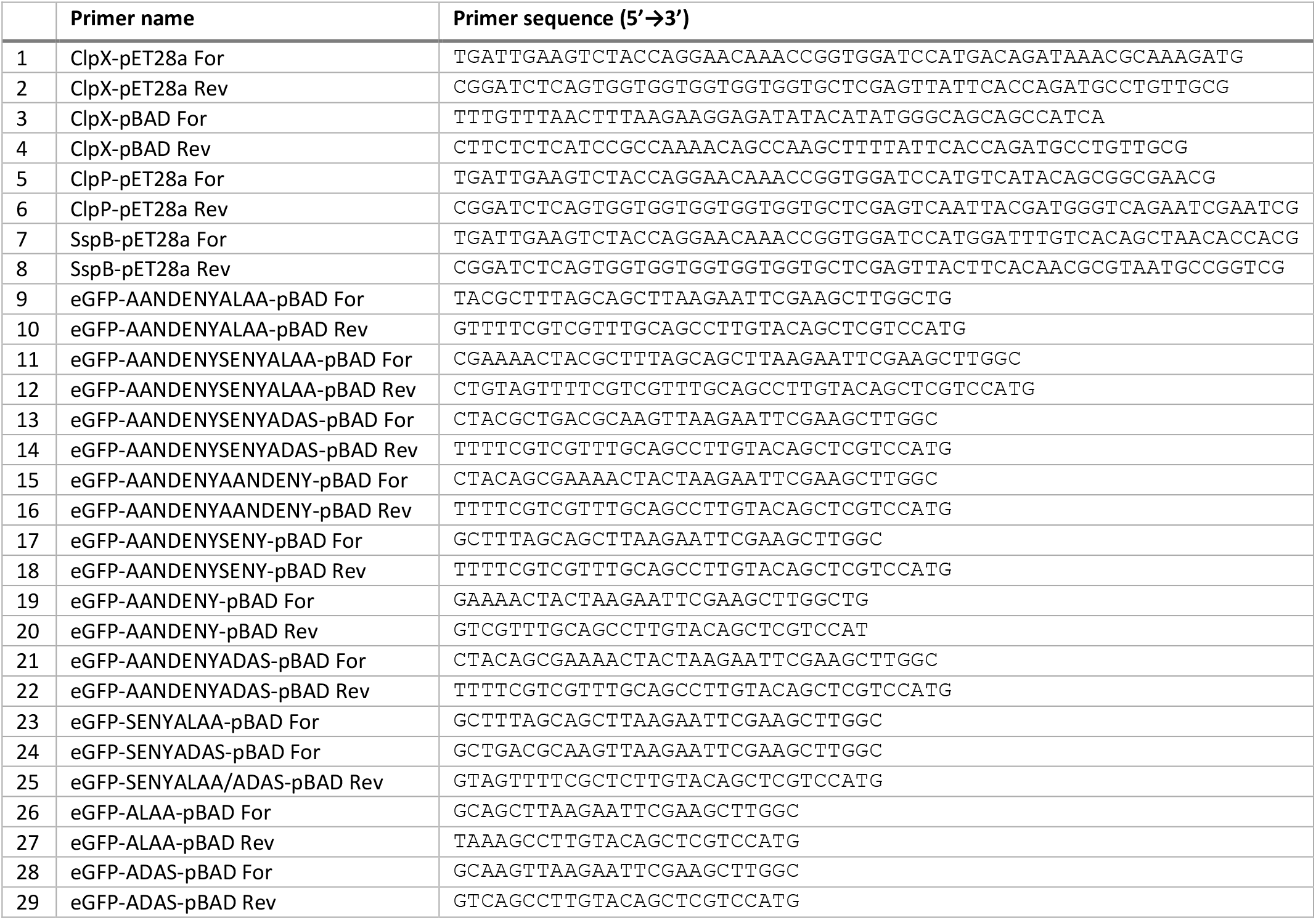
A list of primers used for cloning of the protease complex components and for site-directed mutagenesis of pBAD-eGFP-degron constructs

**Supplementary Table 2.**
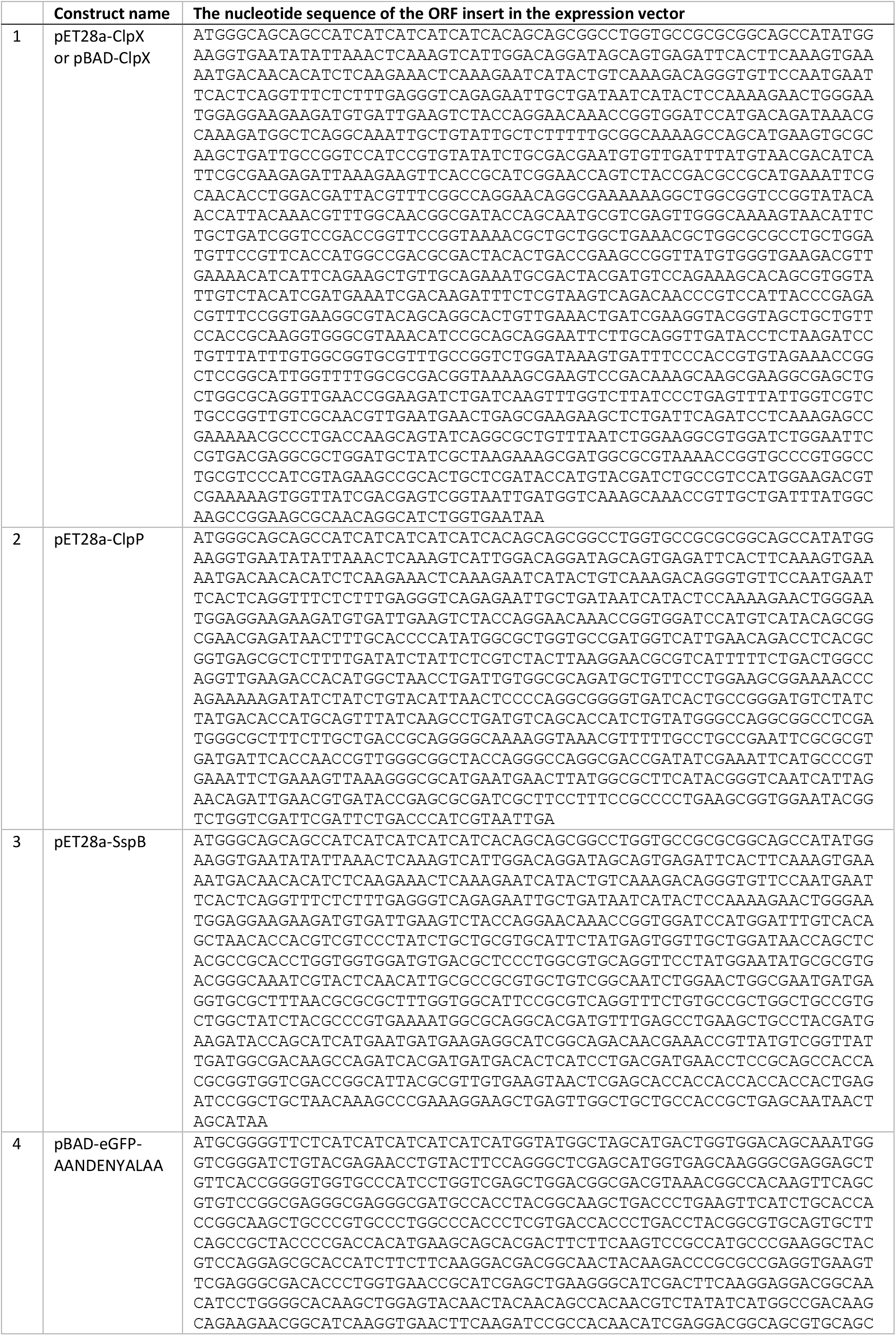
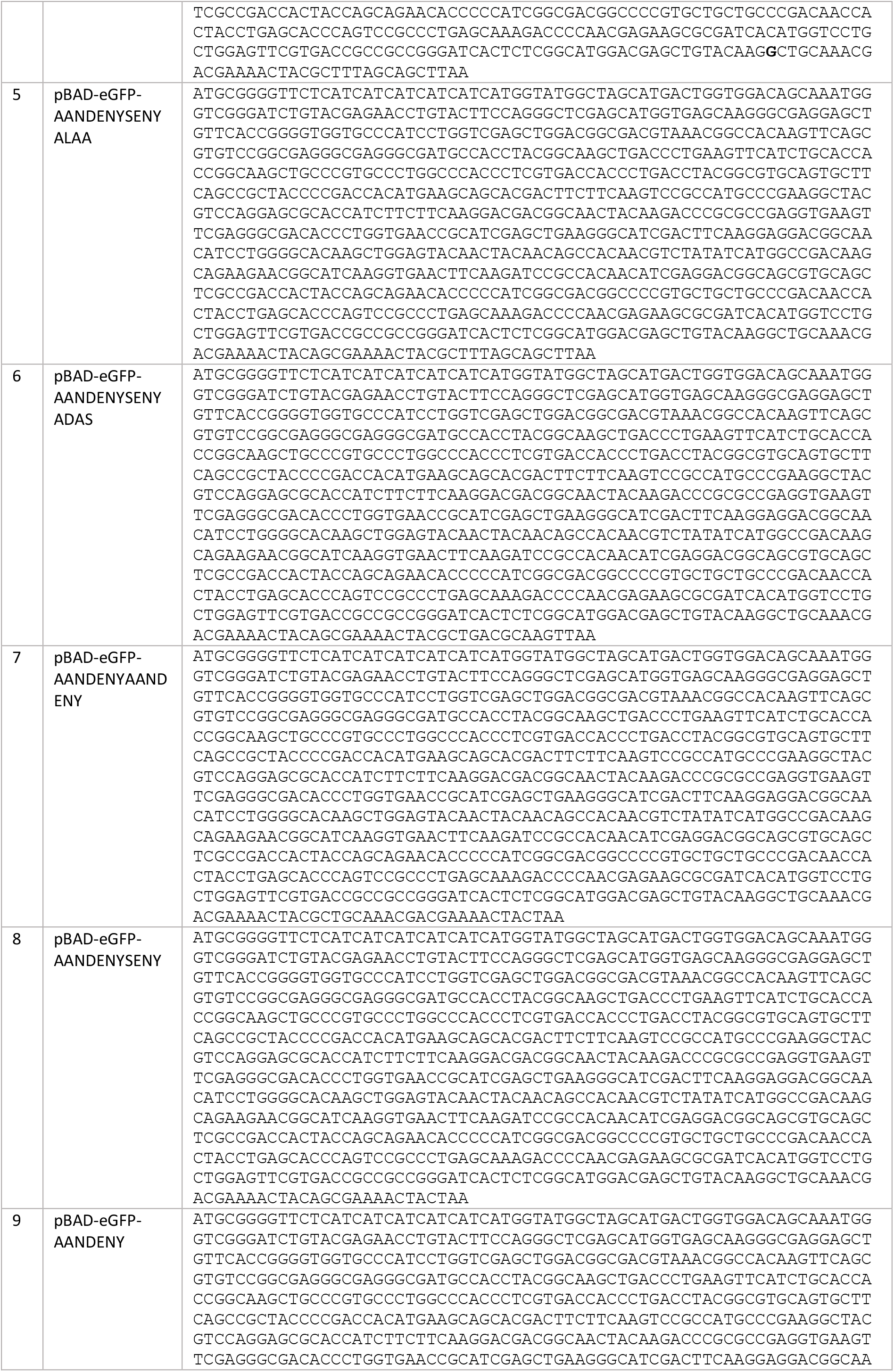
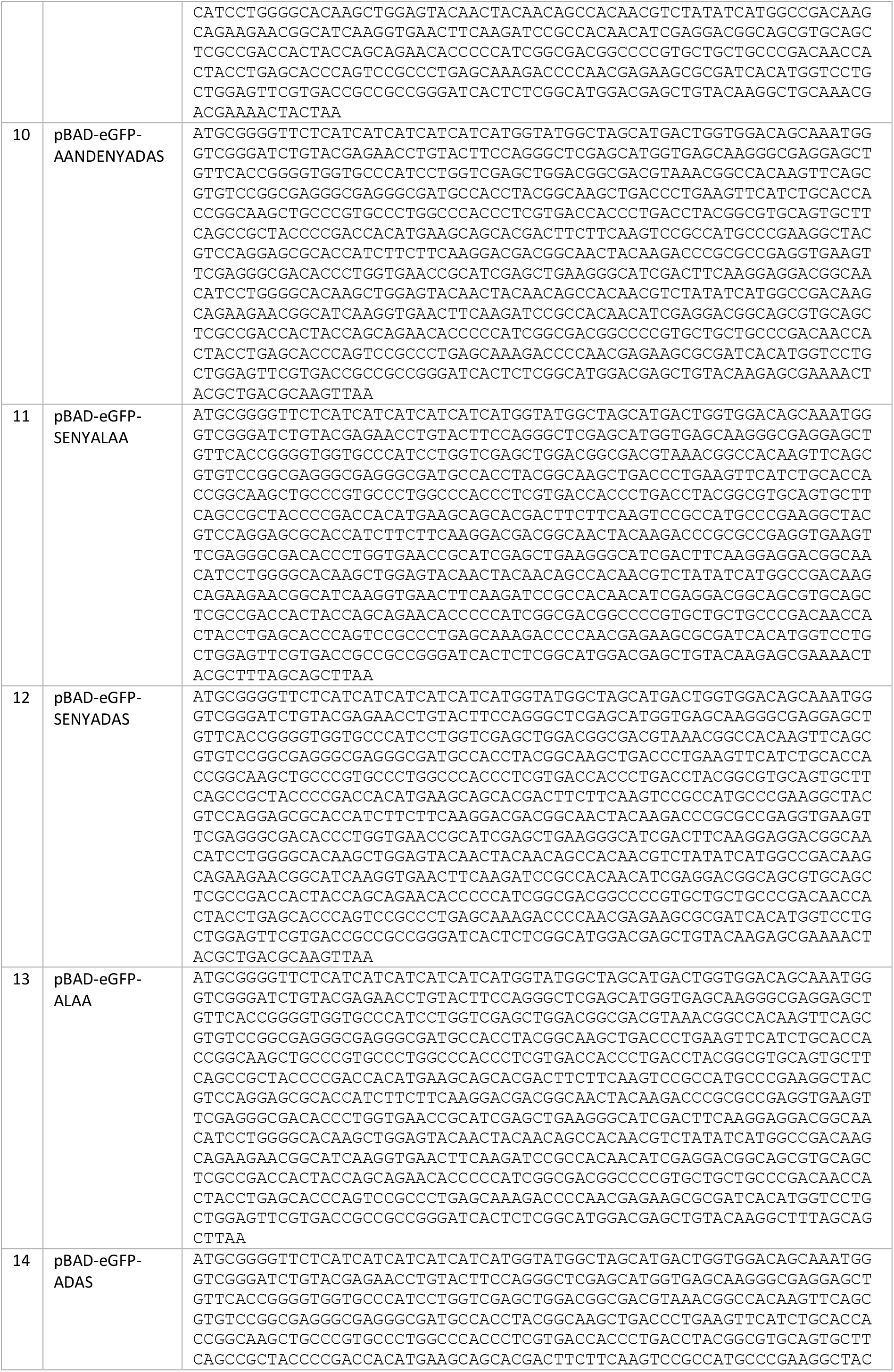
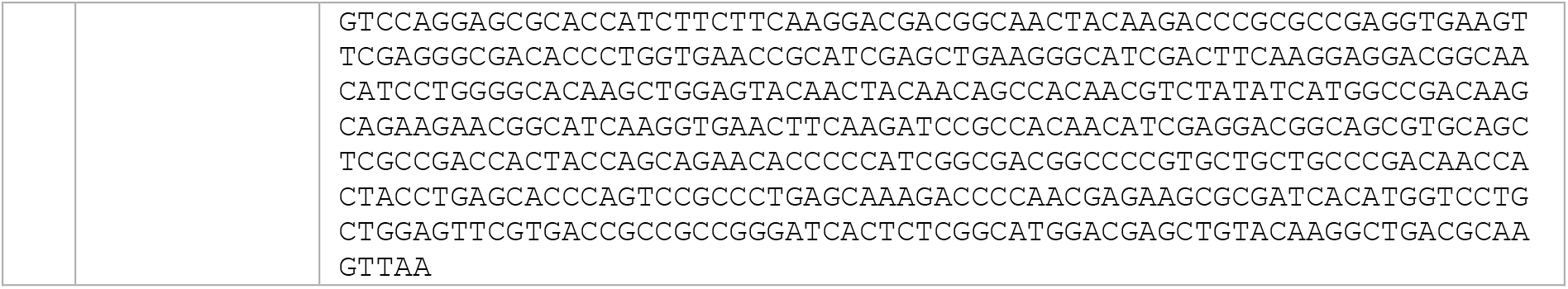
Sequences of the vectors used in the experiments

### SUPPLEMENTARY FIGURES

**Supplementary Figure 1.**
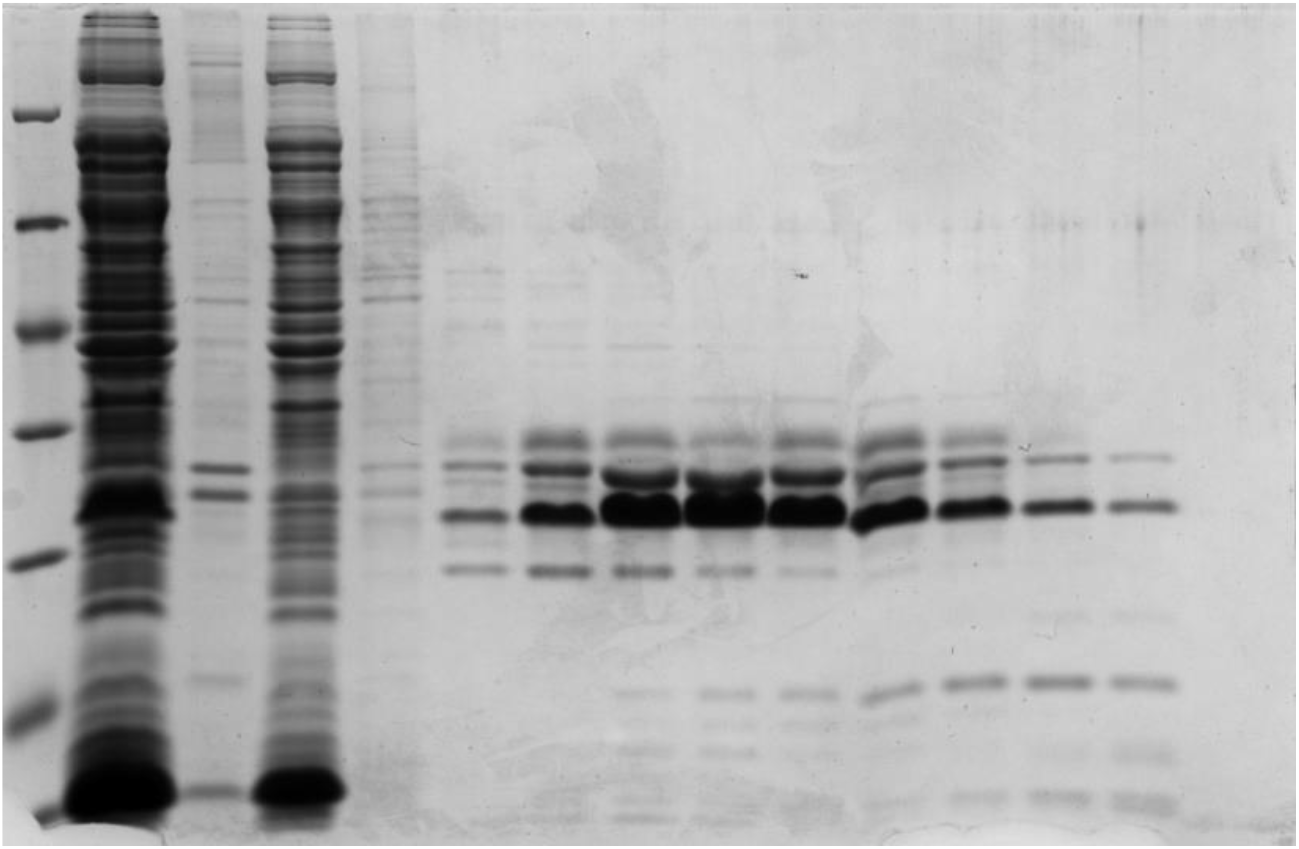
A SDS-PAGE gel with fractions collected after tandem purification of His-TEV-eGFP-AANDENYSENY expressed in *E. coli* Top10

**Supplementary Figure 2.**
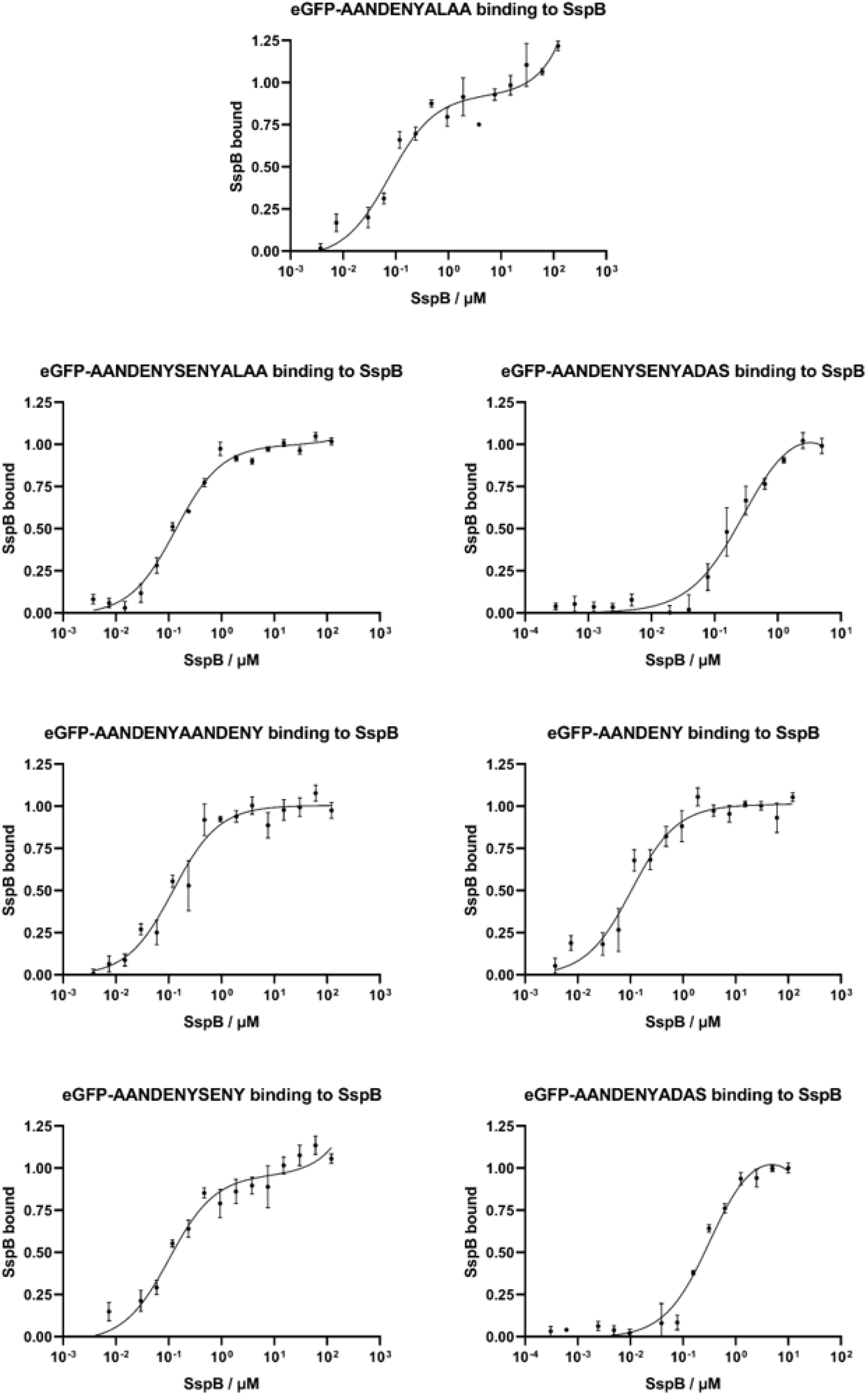
MST results for the binding of eGFP-degrons containing the AANDENY motif to SspB. (a) eGFP-AANDENYALAA; (b) eGFP-AANDENYSENYALAA; (c) eGFP-AANDENYSENYADAS; (d) eGFP-AANDENYAANDENY; (e) eGFP-AANDENY; (f) eGFP-AANDENYSENY; (g) eGFP-AANDENYADAS

